# Geographic variation in evolutionary rescue in a predator-prey system under climate change: an example with aphids and ladybird beetles

**DOI:** 10.1101/2023.03.21.533570

**Authors:** Xuezhen Ge, Jonathan A. Newman, Cortland K. Griswold

## Abstract

Under climate change, species can adapt to changing environments through phenotypic plasticity and natural selection, and this kind of evolutionary adaptation can vary geographically. Most species distribution models (SDMs) are built upon the “Niche conservatism” assumption. They often ignore the possibility of “evolutionary rescue” and underestimate species’ future range limits under climate change. Here, we select aphids and ladybirds as model species and develop an eco-evolutionary model to explore evolutionary rescue in a predator-prey system under climate change. We model the adaptive change of species thermal performance, accounting for biotic interactions of unique life-history trait. Our results show that there is geographic variation in evolutionary rescue for ladybirds (the predator) across different locations in the United States, with ladybirds being more likely to be rescued from extinction in southeastern locations. The possibility of rescue is primarily influenced by the change in seasonality. Our findings also indicate the additive genetic variance of predators has a stronger influence on the phenotype evolution and population dynamics of both prey and predators, compared to the additive genetic variance of the prey. Our research emphasizes the importance of incorporating evolutionary adaptation when predicting species range shift under climate change. The eco-evolutionary model framework can be applied to study the effect of evolution on interacting species’ population abundance and geographic distribution under climate change.

## 1 INTRODUCTION

Over the past two decades, there have been numerous studies on the impacts of climate change on species and how they adapt to these changes (Thomas, 2010; Visser, 2016; Pacifici et al., 2017). Species can adapt to changing environments through phenotypic plasticity and natural selection (Chown et al., 2010; Hoffmann and Sgrò, 2011; van Asch et al., 2013). Climate change has already led to observed shifts in phenologies and physiological capacities, resulting in significant range shifts (Réale et al., 2003; Thomas, 2010; van Asch et al., 2013; Pacifici et al., 2017). Recent reviews have concluded that the effect of evolution on species’ ecological responses to climate change is likely to be more significant for populations at the species’ range margins (Hill et al., 2011; Nadeau and Urban, 2019). This geographic variation emphasizes the importance of considering evolution when predicting species range limits.

Species distribution models (SDMs) have been widely used to how species’ ranges will shift in response to climate change (Elith and Leathwick, 2009; Elith et al., 2010). Many SDM approaches have been developed. These can be classified as correlative SDMs and mechanistic SDMs. Correlative SDMs are the most commonly used approach, but do not directly model the mechanisms behind these relationships (Dormann et al., 2012; Evans et al., 2015). Mechanistic SDMs explicitly define parameters that have a clear ecological interpretation to model biological processes, but may require considerably more effort to collect the detailed biological data, including experimental data (Kearney and Porter, 2009; Evans et al., 2016). Despite the two types of SDMs having their own pros and cons, they both commonly assume that ecological niches are “conserved” when making projections across space or over time (Soberón and Nakamura, 2009). “Niche conservatism” assumes that species tend to retain their fundamental niche over time. This may not be realistic for many species that have been observed to exhibit “rapid evolution”, especially for species with short generations (Ellner et al., 2011; Kopp and Matuszewski, 2014; Garnas, 2018). In range margins, species may be able to persist and avoid extinction through the process of rapid evolution, known as “evolutionary rescue” (Bell, 2017; Nadeau and Urban, 2019). Even if evolution does not fully save the population from extinction, it may still lead to changes in the ecological responses of these species and affect their population sizes and habitat suitability (Meester et al., 2018; Nadeau and Urban, 2019). In addition to the evolutionary adaptation of the target species, there may also be interactions with other as they each evolve in response to climatic changes. There is increasing evidence of the importance of *rapid* evolution on species coexistence for interacting species. Holt et al. (2011) found that predation can strongly affect the evolutionary stability of prey habitat specialization and range limits. Hart et al. (2019) demonstrated that interspecific competition drives rapid genotypic change and causes the population trajectories of the two competing species to converge. Thus, it is crucial to include these processes in SDM projections.

Recently, a variety of model frameworks have emerged to account for species adaptation under climate change and incorporate eco-evolutionary processes into SDMs, such as Allelic adaptive dynamics (ALADYN) (Schiffers and Travis, 2014), RangeShifter (Bocedi et al., 2014), AdaptR(Bush et al., 2016), Gillespie eco-evolutionary models (GEMs) (DeLong and Gibert, 2016), Dynamic eco-evolutionary models (DEEMs) (Cotto et al., 2017), and a macroecological approach (Diniz-Filho et al., 2019). Most of these approaches are individual-based, spatially explicit dynamic models which account for stochastic demographic processes. While these approaches have been successful in considering evolutionary processes to predict species’ range shifts in response to climate change, they also have some important limitations. These approaches may not capture the complexity of real-world trait variation (DeLong and Gibert, 2016). Recent advances in integral projection models (IPMs), such as those developed by Smallegange and Coulson (2013), Rees and Ellner (2016), Janeiro et al. (2017), and Coulson et al. (2021), are able to model evolution under more general conditions, such as non-Gaussian phenotype distribution, as well as stage and age structure.

A number of studies have utilized the ‘moving-optimum model’ to investigate the evolution of quantitative traits (Burger and Lynch, 1995; Matuszewski et al., 2014; Kopp and Matuszewski, 2014). This class of models assume that the optimal values for a particular trait changes over time (Fig. 1a), leading to changes in the distribution of phenotypic values, including a shift in the mean phenotype and a change in phenotypic variance (Fig. 1b). Ectotherms, such as aphids and ladybirds experience changes in their vital rates as the environment moves between optimal and extreme conditions (Sinclair et al., 2016). Thus, these temperature-dependent traits can be represented using Thermal Performance Curves (TPCs), which is captured by thermal optima (*T_opt_*) and thermal tolerances (*CT_min_* and *CT_max_*) (Sinclair et al., 2016). With a moving optimum, the TPCs of a species may evolve over time, potentially impacting the fitness of the species by altering its vital rates (Fig. 1c). The suitability of a habitat for a species may also be influenced by its evolutionary response, potentially leading to a shift in the species’ range in response to climate change. To better understand how species will respond to climate change, it maybe necessary to study how TPCs evolve.

**FIGURE 1.**
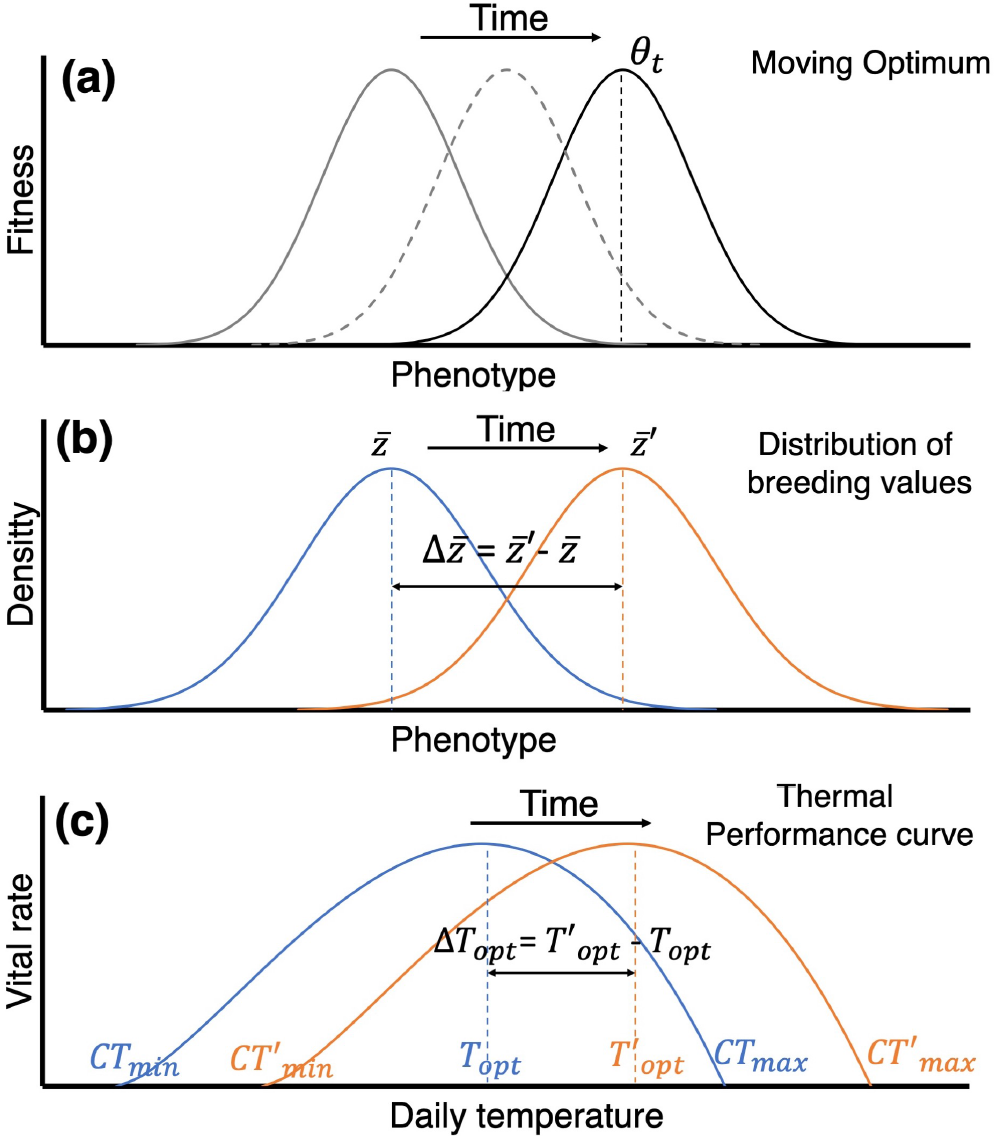
An illustration of trait evolution under climate change and its effect on species range limits under climate change. (a) Solid and dotted curves represent the fitness landscape at different times. *θ_t_* denotes the phenotype with optimal fitness is changing over time. (b) The blue and orange curves represent the distribution of different phenotypes in the population. The mean phenotype evolves from 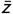 to 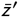 over one generation. (c) The blue and orange curves represent the thermal performance curves for species’ vital rates. The optimum temperature evolves from *T_opt_* to *T_opt_’* with the evolutionary adaptation in response to climate change.

Here, we develop an eco-evolutionary model built on thermal performance. Aphids and ladybirds were chosen as the model species due to the abundant thermal performance data across life history stages. Furthermore, the life histories of aphids and ladybirds are quite different, with one important difference being a shorter generation time for aphids versus ladybirds, but one dominated by asexual (aphids) versus sexual (ladybird) reproduction (Morgan et al., 2001; Parajulee, 2007; Omkar and Pervez, 2005; Raak-van den Berg et al., 2017; Simon et al., 2010). Our model simulates evolutionary adaption with predation of aphids and ladybirds. The reproduction cycle of aphids is complex, because they alternate between sexual and asexual generations (Simon et al., 2010). Without sexual reproduction, independent assortment and recombination could not restore additive genetic variance. Accordingly, the additive genetic variance of aphids may change during the asexual phase. Thus, in contrast to modeling approaches, such as Abrams and Matsuda (1997)‘s predator-prey model, we assume that the additive genetic variance in optimal temperature is constant, but variable for aphids. In addition, we use the integral approach over marginal fitness as defined by Falconer (1960) and Lande (1979), and generalized further by IPM approaches (e.g., Coulson et al., 2021) to capture the change in the distribution of additive genetic effects caused by selection.

To demonstrate the capabilities of our eco-evolutionary model in predicting the range limits of interacting species under climate change, we construct a set of differential equations based on the classic Lotka-Volterra equations (Lotka, 1910), but with an element accounting for the adaptive changes in the optimal temperature (*T_opt_*) for both species. We then select representative geographical locations in the United States at different latitudes and longitudes, and run the model, starting with the locally adapted optimal temperatures (aphid: 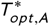; ladybird: 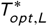) as the initial condition. As part of our analysis, we explore two submodels including one without climatic change and another with a changing climate but no evolution. By comparing the long-term population dynamics of these two submodels with a full model that includes climate change and evolution. We also assess the effect of additive genetic variation on the evolutionary response, as well as compare the roles of climate seasonality and warming trend on ecological outcomes. Our results demonstrate the effect of a moving *T_opt_* on species habitat suitability and highlight the complex interplay between climate change, evolution and geography.

## 2 METHODS

### 2.1 Biology of the aphid and ladybird

Aphids and ladybirds are a good model system to study the interplay between predation, climate change and adaptation when generations are short (Tayeh et al., 2015; Simon and Peccoud, 2018). There are numerous experimental studies of thermal performance on these species, which can provide empirical support for setting model parameters (Morgan et al., 2001; Parajulee, 2007; Omkar and Pervez, 2005; Raak-van den Berg et al., 2017). In this study, we are using the general biological characteristics of aphids and ladybirds rather than the specific characteristics of a single species as we are not focused on studying the evolutionary effects of a particular pair of species. As noted in the introduction, we assume in a geographic location, aphids and ladybirds are initially adapted to these environment.

Aphids and ladybirds have distinct life cycles and reproduction strategies. The life cycle of aphids include both asexual and sexual generations. Aphids primarily reproduce many generations asexually during most of the year, and reproduce sexually once during the autumn (Dixon, 1977). For many aphid species, the occurrence of a sexual generation is triggered by environmental cues such as short-day length and low temperatures (Ogawa and Miura, 2014). Ladybirds, on the other hand, only reproduce sexually and have fewer generations per year than aphids (Pervez and Omkar, 2006).

The vital rates for both aphids and ladybirds are sensitive to environmental temperatures (Hullé et al., 2010; Raak-van den Berg et al., 2017). The shape parameters for the TPCs of these vital rates vary depending on the species and their developmental stages (Zhao et al., 2017; Jalali et al., 2014). For simplicity, we assume aphid and ladybird have the same functional shape (*q*_1_ and *q*_2_ in Eq. 12, *k*_1_ and *k*_2_ in Eq. 13) to their TPCs. In our eco-evoultionary model, we assume the TPCs may shift horizontally with climate change, meaning that the three TPC parameters *CT_min_*, *CT_max_* and *T_opt_* evolve together at the same rate and direction. Thus, we substitute *CT_min_* and *CT_max_* with functions of *T_opt_* to keep the evolutionary component concise. We use non-stage structured models for both aphids and ladybirds, such that we assume progression across life history stages is captured by single parameters for both species. Figure 2 illustrates the reproduction phrases and their corresponding evolving traits for aphids and ladybirds.

**FIGURE 2.**
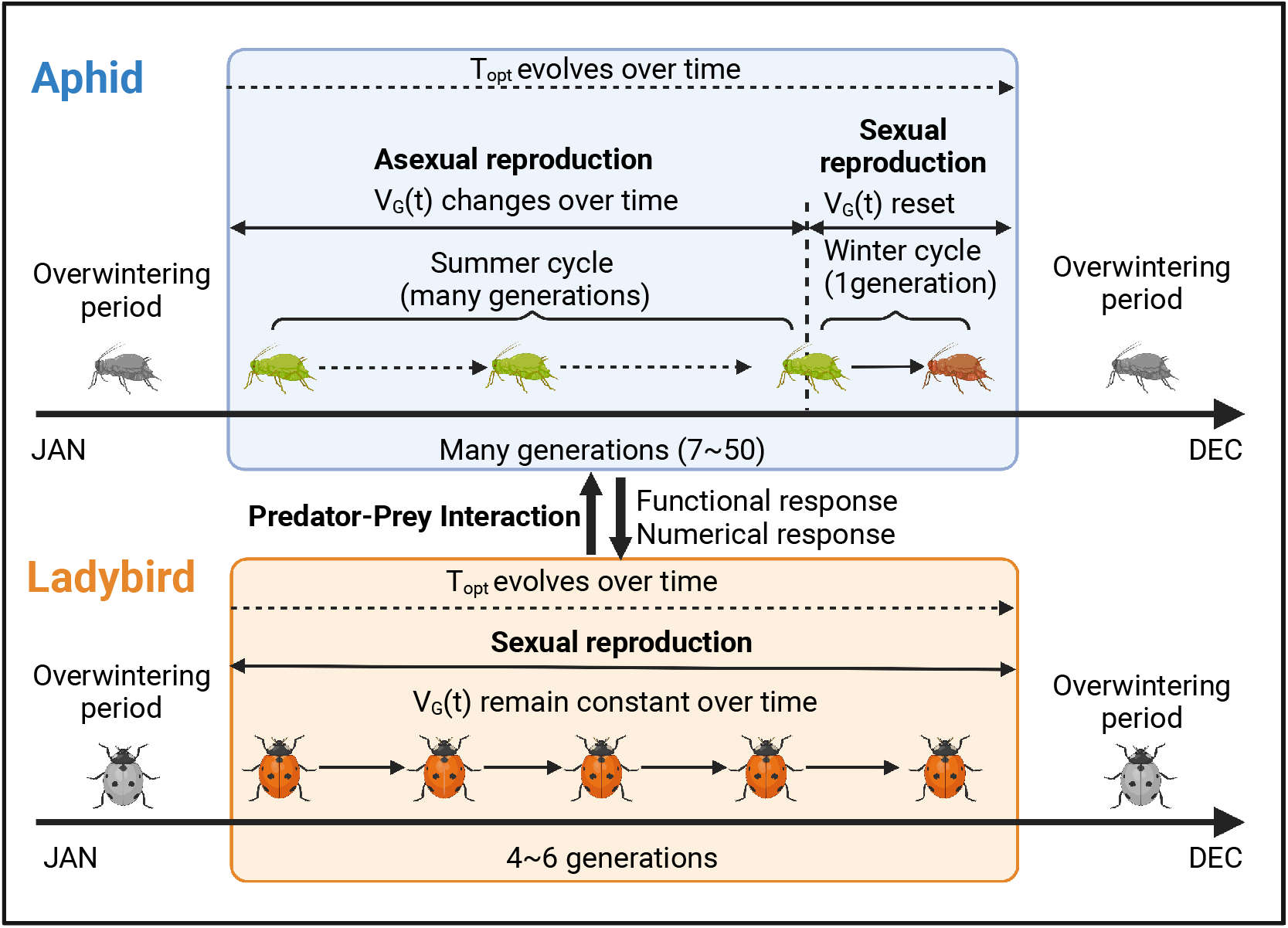
Schematic diagram of the life cycle for the aphid and ladybird within a year. Aphids reproduce asexually for most of the year and sexually once in the autumn. Both the additive genetic variance (*V_G_*) and optimum temperature (*T_opt_*) evolve over time during the asexual phase, but *V_G_* is assumed to reset during the sexual phase by meiotic segregation, recombination and random mating. Ladybugs only reproduce sexually, and we assume remains constant and *T_opt_* evolves over time.

### 2.2 Climate data for selected geographical locations

To examine geographic variation of the evolutionary response, we selected nine geographical locations from the agricultural regions in the United States (Table 1). These nine locations are evenly distributed across the area between 32°N – 44°N, and 84°W – 104°W.

**TABLE 1.** Locations and temperature parameters of the nine geographical locations in the United States. Seasonality is half the difference between the yearly temperature minimum and maximum at a given location. The trend is the change in mean temperature over 100 years. These projections come from the French CNRM-CM6-1 model using the highest emission scenario (i.e., SSP5-8.5; Coupled Model Intercomparison Project Phase 6 (CMIP6) (Voldoire, 2018). For more information see http://www.umr-cnrm.fr/cmip6/spip.php?article11.

| ID | Latitude | Lognitude | Location | Tmean ($\bar{T}$)<br>°C/year | Seasonality ($s$)<br>°C/year | Trend ($k$)<br>°C/100 years |
| --- | --- | --- | --- | --- | --- | --- |
| SW | 32°N | 104°W | Angeles, Texas | 13.318 | 12.398 | 7.814 |
| SO | 32°N | 94°W | Logansport, Louisiana | 16.889 | 10.904 | 5.779 |
| SE | 32°N | 84°W | Cobb, Georgia | 15.973 | 9.942 | 5.793 |
| CW | 38°N | 104°W | Fowler, Colorado | 5.055 | 13.699 | 8.109 |
| CO | 38°N | 94°W | Taberville, Missouri | 11.252 | 15.805 | 7.138 |
| CE | 38°N | 84°W | Mt. Sterling, Kentucky | 10.723 | 13.264 | 6.476 |
| NW | 44°N | 104°W | West Pennington, South Dakota | 4.153 | 16.159 | 7.734 |
| NO | 44°N | 94°W | Beauford, Minnesota | 5.103 | 19.705 | 8.290 |
| NE | 44°N | 84°W | Standish,Michigan | 6.477 | 15.917 | 7.579 |

We downloaded the future daily temperature data from a Gobal Circulation Model which has a relatively higher spatial resolution (i.e., CNRM-CM6-1) and selected the highest emission scenario (i.e., SSP5-8.5) to obtain the future climate projections with the most extreme global warming trend (Coupled Model Intercomparison Project Phase 6 (CMIP6) (Voldoire, 2018). We generated “smoothed” temperature data using spectral decomposition of the daily temperature time series data into three components (trend, seasonality, and noise) and removing the “noise” component. The remaining two components, which form our “smoothed” temperature data, contain the temperature seasonality and the global warming trend. We used “smoothed” temperature data to eliminate the effect of day-to-day variability; accordingly our TPC model assumes an inherent buffering against day-to-day variability.

Similar to Ge et al. (2022), we use a cosine function of daily mean temperature to generate temperature profiles for each of the geographical locations (Eq. 1). In Eq. 1, *t* denotes time (day), 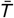 denotes the annual mean temperature (Eq. 2), *k* denotes the trend of temperature change every century, and *s* denotes the ‘seasonality’ of daily temperature, which is defined as half of the difference between yearly minimum temperature (*T_min,year_*) and yearly maximum temperature (*T_max,year_*) in a year (Eq. 3).

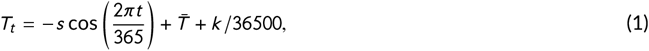

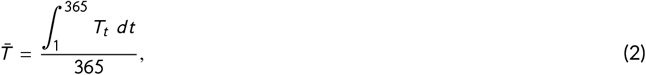

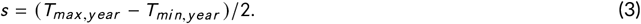

According to the definitions of the three temperature parameters 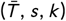, we fit a linear regression to the “trend” component of the decomposed time series temperature data and estimated 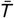 and *k*, then we calculated *s* by estimating amplitude of the “seasonality” component. The temperature parameter values of the nine locations are listed in Table 1. With these temperature parameters, we generated the daily temperature data from 2000 to 2150 at each location for model simulations.

### 2.3 Model description

#### 2.3.1 Genetic adaptation

In this study, we simulate the population dynamics of predators (i.e., ladybirds) and prey (i.e., aphids), with their population densities denoted as *L* and *A,* respectively. The predator and prey populations are both assumed to have a moving trait (*T_opt_*), which demonstrates species’ genetic adaptation in thermal performances in response to a warming climate. Both species’ vital rates are functions of air temperature (T) and mean values of *T_opt_* (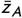 or 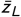). The model equations, which include temperature-dependent and evolving vital rates, are

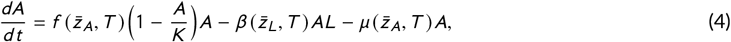

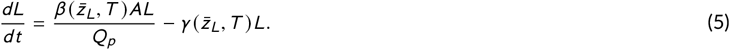

In Eq. 4 and Eq. 5, *K* is the carrying capacity for aphid population. 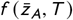 represents the temperature-dependent growth rate for aphids. 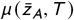 and 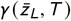 represent the intrinsic mortality rates for aphids and ladybirds, respectively. For the prey, we assume that predation by predators is the only source of extrinsic mortality. For the predator, we assume that predator birth rates depend on prey capture rates, and that the only source of extrinsic mortality in our model derives from starvation at low prey population sizes. Predator-prey interactions are determined by the functional and numerical responses of ladybirds. Previous research has suggested that a Type II functional response is commonly observed in ladybirds (Zarghami et al., 2016; Sharma et al., 2017). Thus, we use Holling’s (1959) Type II functional response to describe the relationship between the number of consumed prey and the prey density, and the function 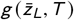 to regulate the effect of temperature on the functional response. Together, the rate for each aphid is predated by each ladybird 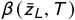 is

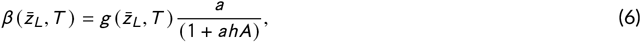

where *a* and *h* denote predator’s searching rate and handling time, respectively (Holling, 1959). The predator’s numerical response is based on the assumption that reproduction rate of predators is proportional to the number of prey consumed (Solomon, 1949). Thus, we use transformation rate (denoted by *Q_p_* in Eq. 5) to represent the the mean number of aphids a ladybird needs to consume to reproduce a single egg. All together, the number of consumed prey contributed to population growth of predators is 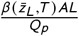. All the parameter values for these vital rates are listed in Appendix S1 and Table S1.

From equations 4-6, the intrinsic population growth rate (continuous growth rate) for the predator and prey population is

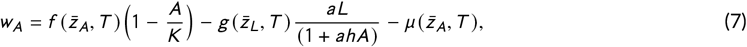

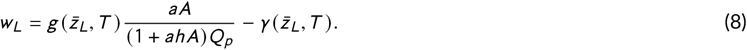

The discrete growth rates for the aphid and ladybird are exp (*w_A_*) and exp(*w_L_*) over a discrete time step. At a discrete time step in the numerical solution of the system defined by equations 4 and 5, there is initially a mean breeding value 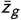, and additive genetic variance *V_G_*, as well as environmental variance *V_E_* (Note that we use *V_G_* in this study to represent additive genetic variance because we use letter ‘A’ to represent aphid). We assume breeding values and environmental effects are normally distributed. According to the integral approach defined by Falconer (1960) and Lande (1979), and generalized further by Coulson et al. (2021), the distribution of breading values *P*(*z_g_*) following selection is:

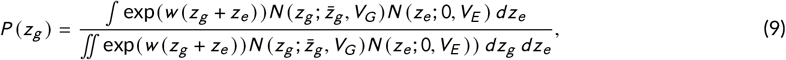

where 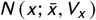 is the normal distribution of the random variable *x* with the mean *x* and variance *V_x_*. The mean breeding value after selection 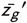 and additive genetic variance 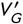 are then,

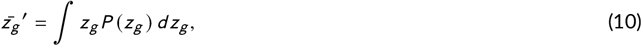

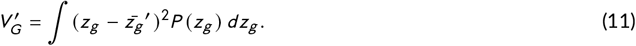

Although selection can cause small deviations from normality, in aphids we assume in the next time step breeding values are normarlly distributed with variance 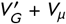 (See next section). In ladybirds selection can cause a small change in *V_G_*, but we assume continuous sexual reproduction restore to a baseline level.

#### 2.3.2 Generalized equations for vital rates

In our models, we mainly use the following two generalized equations (Ge et al., 2022) to model the effect of temperature on species vital rates (i.e., 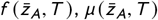 and 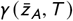) and predation rate 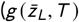, albeit with differing values of the shape parameters:

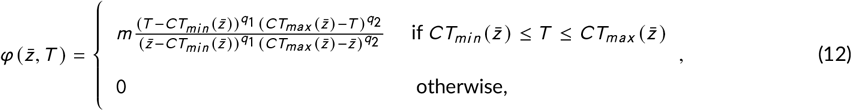

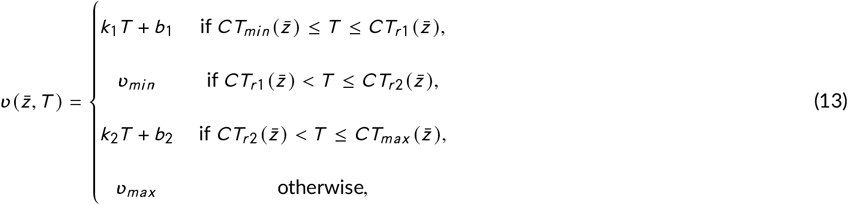

such that, for example, 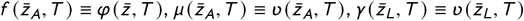.

In Eq. 12 and Eq. 13, *T* is the air temperature. *CT_min_* and *CT_max_* are the minimum and maximum temperature thresholds beyond which growth rate is nil, and mortality rate is maximal. 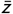 is the mean phenotypic value of *T_opt_*, which the optimum temperature at which the vital rate reaches to its maximum (m). *q*_1_ and *q*_2_ are shape parameters which adjust the skewness of the thermal performance curve. *CT*_*r*1_ and *CT*_*r*2_ define the temperature range in which mortality is minimal. *k*_1_, *b*_1_, *k*_2_ and *b*_2_ are the parameters which make *ν*(*CT*_*r*1_) = *ν*(*CT*_*r2*_) = *ν_max_* and *ν*(*CT_min_*) = *ν*(*CT_max_*) = *ν_min_*. We assume the shape of species’ thermal performance curves are always the same no matter how the habits differs among different geographical locations or changes under future climate conditions, and only shift horizontally. Therefore, all the thermal performance traits (e.g., *CT_min_*, *CT*_*r*1_) can be measured by a single parameter (*T_opt_*). For all the fixed thermal performance curves, *q*_1_ and *q*_2_ are fixed values which are *q*_1_ = 1.5 and *q*_2_ = 1. All the parameter values for Eq. 12 and 13 are listed in Appendix S1 and Table S1.

#### 2.3.3 Model sets

We construct three models to investigate the effects of climate change, and evolutionary adaptation on the population dynamic of aphids and ladybirds. By decomposing the individual contributions of each of these factors, we are able to gain a better understanding of their effects on the predator-prey system and the complex interactions between these factors.

##### Model 1: No evolution and a constant climate

The first model we construct simulates a constant climate for aphids where the daily temperatures have an annual pattern, but the pattern doesn’t change through time (i.e., no interannual variation). Thus, the aphids and ladybird do not experience the effects of climate change. We find values 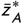 and 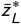, such that they are at equilibrium. All of the rates described in Eq. 4 and Eq. 5 are temperature-dependent (Eq. 12 and Eq. 13) but have the same annual patterns in each year. This model provides a baseline to identify the predator and prey dynamics in a constant environment, and can also be used to compare the effects of climate change and evolution on these dynamics.

##### Model 2: No evolution, but with climate change

In this modeling scenario, we assume aphids and ladybirds are living in a warming environment, unlike the constant environment assumed in the baseline model. However, their thermal performances are notable to adapt evolutionarily to the changing conditions, i.e., *V_G_* for both aphids and ladybirds is zero. As a result, the mean phenotypic values (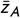 and 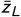) remain constant. Despite this, the annual patterns of these rates will still be affected by the interannual variation in temperature data, which alters the annual population dynamics of aphids and ladybirds.

##### Model 3: Eco-evolutionary model with climate change

In the eco-evolutionary model, we incorporate species’ evolutionary adaption by allowing 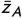 and 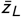 to change through time in response to climatic change, i.e., *V_G_* > 0 for aphids and ladybirds. The changes in breeding value and additive genetic variance, and population densities for aphids and ladybirds due to nature selection are

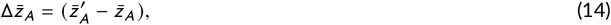

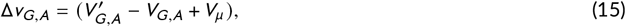

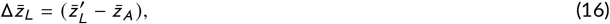

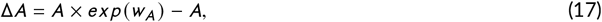

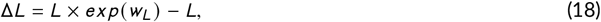

where *w_A_* and *w_L_* represents the intrinsic rates of increase for aphids and ladybirds, which are given by Eq. 7 and Eq. 8. In Eq. 15, *V_μ_* represents the variance in mutational effect and is assumed equal to 0.001 × *V_E_* (Lynch and Walsh, 1998). According to the experiments conducted by Logan et al. in 2020, we set the values of *V_G,A_* = *V_G,L_* =0.3, *V_E,A_* = *V_E,L_* =0.7 based on the estimated additive genetic variance and heritability for the thermal performance of the harlequin beetle (*Harmonia axyridis).*

### 2.4 Model simulations

#### 2.4.1 Locally adapted *T_opt_* for aphids and ladybirds

To model the population dynamics of aphids and ladybirds in various geographical locations, we use daily temperature data in 2000 to estimate the locally adapted optimal temperature for each species (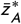 for aphids and 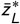 for ladybirds). These estimates are used as the starting values for our long-term simulations. We follow the procedures below to estimate 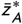 and 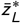.

To begin, we obtain the maximum daily temperature data in 2000 (*T*_*max*,2000_) and used its floor value (i.e., the greatest integer that is less than or equal to *T*_*max*,2000_) as the initial values for 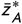 and 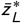.

Next, we run the eco-evolutionary model (Model 3) for a period of five years using daily temperature data in 2000. By analyzing the output value of daily evolving 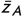 through a linear regression, we are able to estimate the trend of 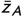 and 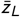. If the trend value is positive, it means that the value of 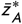 is too low and needs to be increased. Conversely, a negative trend value indicates that 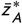 is too high and needs to be decreased. To determine the optimal value of 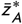, we continually adjust its value by 0.5 °C until the sign of the trend changes.

Once we determine the locally adapted optimal temperature for aphids 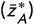, we repeat the above procedures to estimate the local adapted optimal temperature for ladybirds 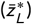. Based on 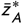 and 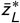, we can calculate the other thermal performance parameters and obtain the thermal performance traits that best match the local temperature data in 2000 for each species. The estimated values of 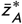 and 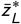 for each geographical location are shown in Fig. 4.

#### 2.4.2 Long-term evolutionary effects on population dynamics

For each geographical location, we use temperature profiles specific to that location and the locally adapted optimal temperatures (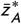 and 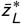) for aphids and ladybirds to run the three models described in section 2.3.3 from 2000 to 2150.

We set the starting conditions to be same for each model (*A* = 1000000, *L* = 5000, 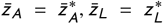). We account for the overwintering periods for both species. At the beginning of each year, we introduce the aphids and ladybirds into the model when the temperature is warm enough to support positive aphid and ladybird population growth rates (*w_A_* > 0 or *w_L_* > 0), respectively. Aphids switch from their asexual phase to their sexual phase when the climate becomes cooler and daylength becomes shorter (Ogawa and Miura, 2014). For simplicity, we use the date when aphids switch phases as the date when both species enter overwintering periods. During the overwintering period, aphids and ladybirds will terminate their population growth and pass through the cold winter. Due to lack of empirical data, we arbitrarily assume the year to year overwintering mortality rate for both species is 0.2. Daylength (*DL*) represents the required number of daylight hours and is calculated as follow (Langille et al., 2016):

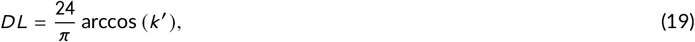

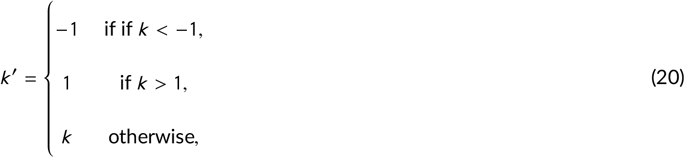

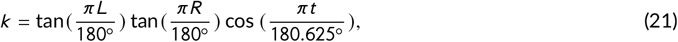

where *DL* is determined by *k*, which is the exposed radius between the sun’s zenith and the sun’s solar circle and estimated by a function of latitude, L, the number of days from January 1st, *t* and the Earth’s rotational axis, R = 23.439° (Langille et al., 2016). Then, we use *s* to act as a computational control “switch” to enable the sexual phase, aphids remain in their asexual phase when *s* = 0 and switch to their sexual phase when *s* = 1:

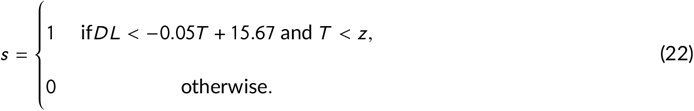

After both species enter the overwintering period: (*i*) both species stop their development, (*ii*) the mean phenotypes for aphid and ladybird remain unchanged until they emerge in the next year, and (*iii*) aphid additive genetic variance is restored to its initial value due to the sexual reproduction. Aphid and ladybird population size will reduce 20% due to the overwintering mortality rate and the remaining aphids and ladybirds (80%) will be used as the start values in the next year.

To study the long-term evolution of the aphids and ladybirds, we first run the eco-evolutionary model from 2000 to 2020 to flush the initial conditions and begin the simulation from 2021 with realistic starting values. We then run the three models separately from 2021 to 2150 to study how evolution changes aphid and ladybird populations over a period of 130 years. The first model uses the daily temperatures in 2020 to simulate a constant environment, while the second and third models use the daily temperatures from 2021 to 2150, which includes the effects of climate change. The second model doesn’t not consider evolution by setting both species’ additive genetic variances to be 0, while the third model takes into account evolution by setting *V_G_ >* 0. Based on the output of daily population abundance for aphids and ladybirds from three models, we calculate the accumulation of the daily population abundance within each year and denote them as “annual aphid pressure” (*AAP*), and “annual ladybird pressure” (ALP). For brevity, we will refer them as aphid and ladybird population abundances throughout the paper. By comparing the long-term patterns of aphids and ladybirds’ population abundances in all three models, we can examine the impact of evolutionary adaptation on each species.

#### 2.4.3 Effects of additive genetic variance and heritability

As noted by Lande (1979, eq. 5b), a general expression for the response in phenotype (*z*) across a generation is:

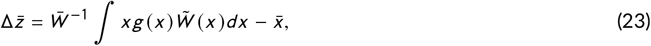

where 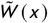 is the marginal fitness of an individual with breading value *x*, 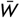 is mean fitness of the population and *g*(*x*) is the distribution of breeding values. Lande (1979) applied the gradient to a multivariate form of Eq. 23 to obtain the multivariate breeder’s equation, which in univariate form is:

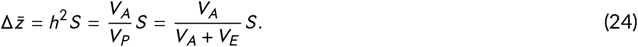

Here, *s* is the selection differential which is the difference in mean phenotype before and after selection. *h*^1^ is the narrow-sense heritability, *V_A_* and *V_E_* are the variances explained by additive genetic effects and environmental effects, *V_P_* is the total variance of the trait in the population. For a given phenotype *z*, Eq. 24 implies that the response to selection increases with increasing *VA* because both *h*^2^ and *s* will increase. The effect of *V_E_* may have unpredictable consequences because increasing *V_E_* leads an increasing *S* but a decreasing *h*^2^. It’s not straightforward to determine whether the increase in selection differential surpasses the decrease in heritability.

We did a simulation to test the effect of *V_E_* with varying predator’s transformation rate *Q_p_*. Fig. 3 presents marginal fitness as a function of the breeding value of *T_opt_* when these breeding values are integrated over normally distributed environmental effects, the thermal performance function and absolute fitness. We used Lande’ integral and fitness gradient approach and got equal responses of selection. Both of approaches indicate the shape of marginal fitness landscape varies for different predator species, which may yield different responses to selection. A predator species that requires more prey to reproduce an offspring (higher transformation rate, *Q_p_*) has stronger responses to selection with increasing *V_E_*, indicating that the increase in selection differential surpasses the decrease in heritability as environmental variance increases. A predator species with a lower transformation rate could have an opposite change in responses to selection with increasing *V_E_*. Our analysis indicates the uncertainty of environmental effect on response to selection, thus we mainly focus on the effect of genetic variance in this study.

**FIGURE 3.**
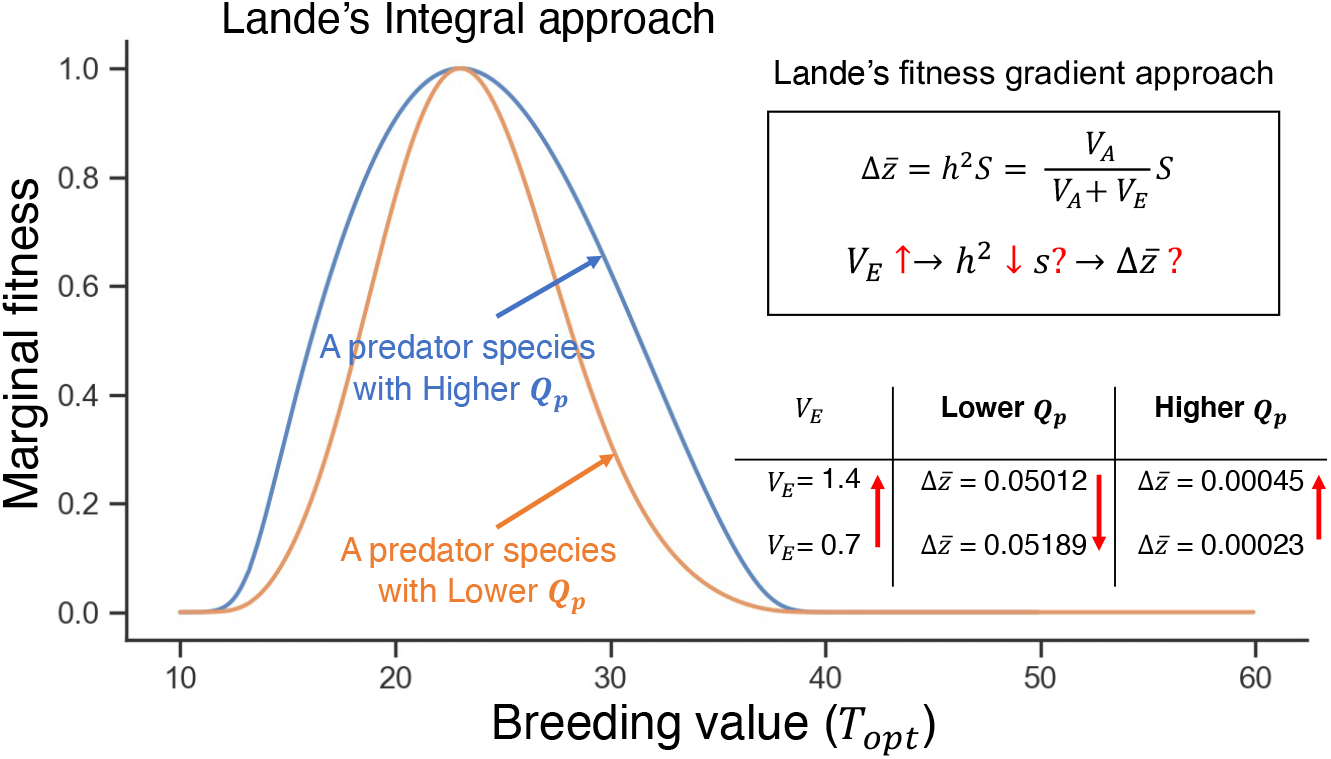
Illustration of marginal fitness landscapes and response to selection estimated by Lande’s integral and fitness gradient approaches. *Q_p_* represents the transformation rate, i.e., the number of aphids a ladybird needs to consume to reproduce a single egg. The blue and orange curves represent the shapes of the marginal fitness landscape when using the integral approach. As we increase the value of *Q_p_*, the shape of marginal fitness landscape changes, as well as the effect of an increase in *V_E_*.

To understand how populations may evolve in response to climate change within a predator-prey system, we adjust the values of additive genetic variance in each species, and use these values to construct various scenarios with different evolutionary potential. This can provide insight into the potential for populations to evolve rapidly in the face of climate change.

To determine the individual effects of *V_G,A_* and *V_G,L_*. we manipulate their baseline values by a factor of ten. We also set either *V_G,A_* or *V_G,L_* to be 0 to to simulate a scenario in which only prey or predator is evolving (Table 2). Since narrow-sense heritability is given by 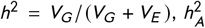 and 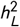 will then vary in proportion to the changes in *V_G,A_* and *V_G,L_*. Among the nine geographical locations, we selected one southern (Logansport, Louisiana) and one northern (Beauford, Minnesota) location and ran the eco-evolutionary model for 100 years. By studying the long term population dynamics of aphids and ladybirds under different sets of genetic parameters, we assess the influence of additive genetic variance and heritability on a species’ evolutionary potential to adapt to climate change.

**TABLE 2.** Different sets of parameters for estimating the effects of additive genetic variance(*V_G_*) and heritability *h*^2^. The baseline is set according to the experiments conducted on the thermal performance of the harlequin beetle (*Harmonia axyridis*) by Logan et al. in 2020.

| Parameters set | $V_{G,A}$ | $V_{G,L}$ | $V_{E,A}$ | $V_{E,L}$ | $h_A^2$ | $h_L^2$ |
| --- | --- | --- | --- | --- | --- | --- |
| **Baseline** | **0.30** | **0.30** | **0.70** | **0.70** | **0.30** | **0.30** |
| $V_{G,A}$-Low | 0.03 | 0.30 | 0.70 | 0.70 | 0.04 | 0.30 |
| $V_{G,A}$-High | 3.00 | 0.30 | 0.70 | 0.70 | 0.81 | 0.30 |
| $V_{G,L}$-Low | 0.30 | 0.03 | 0.70 | 0.70 | 0.30 | 0.04 |
| $V_{G,L}$-High | 0.30 | 3.00 | 0.70 | 0.70 | 0.30 | 0.81 |
| $V_{G,A}$-0 | 0.00 | 0.30 | 0.70 | 0.70 | 0.00 | 0.30 |
| $V_{G,L}$-0 | 0.30 | 0.00 | 0.70 | 0.70 | 0.30 | 0.00 |
| $V_{G,A}$-$V_{G,L}$-0 | 0.00 | 0.00 | 0.70 | 0.70 | 0.00 | 0.00 |

## 3 RESULTS

### 3.1 Geographic variation in evolutionary rescue

Figure 4 shows annual population dynamics for the aphid and ladybird across the nine geographical locations. For the three model scenarios, aphid and ladybirds population have different dynamics over 150 years. In each subplot of Fig. 4a-b, the black curve shows the annual population abundances of locally adapted aphids and ladybirds over time in a constant climate (Model 1). Both species are able to maintain their populations and eventually reach a stable equilibrium. In Model 2, which takes into account climatic change but assumes no evolutionary response (red dashed curves) and locally adapted starting conditions, the population of aphids declines, while the population of ladybugs goes extinct due to their inability to adapt to the changing climate conditions and reduced prey availability. In the eco-evolutionary model (Model 3, shown by the red solid curves in Fig. 4a-b), we found that the mean optimal thermal performance (*T_opt_*) of both species increases as a result of natural selection. This adaptation allows the aphid population to persist and maintain its population size (Fig. 4a), and the ladybird population is rescued from extinction in some southeastern locations. In northwestern locations, the ladybird population reached a relatively low equilibrium, which is not visible in the subplots due to scaling.

**FIGURE 4.**
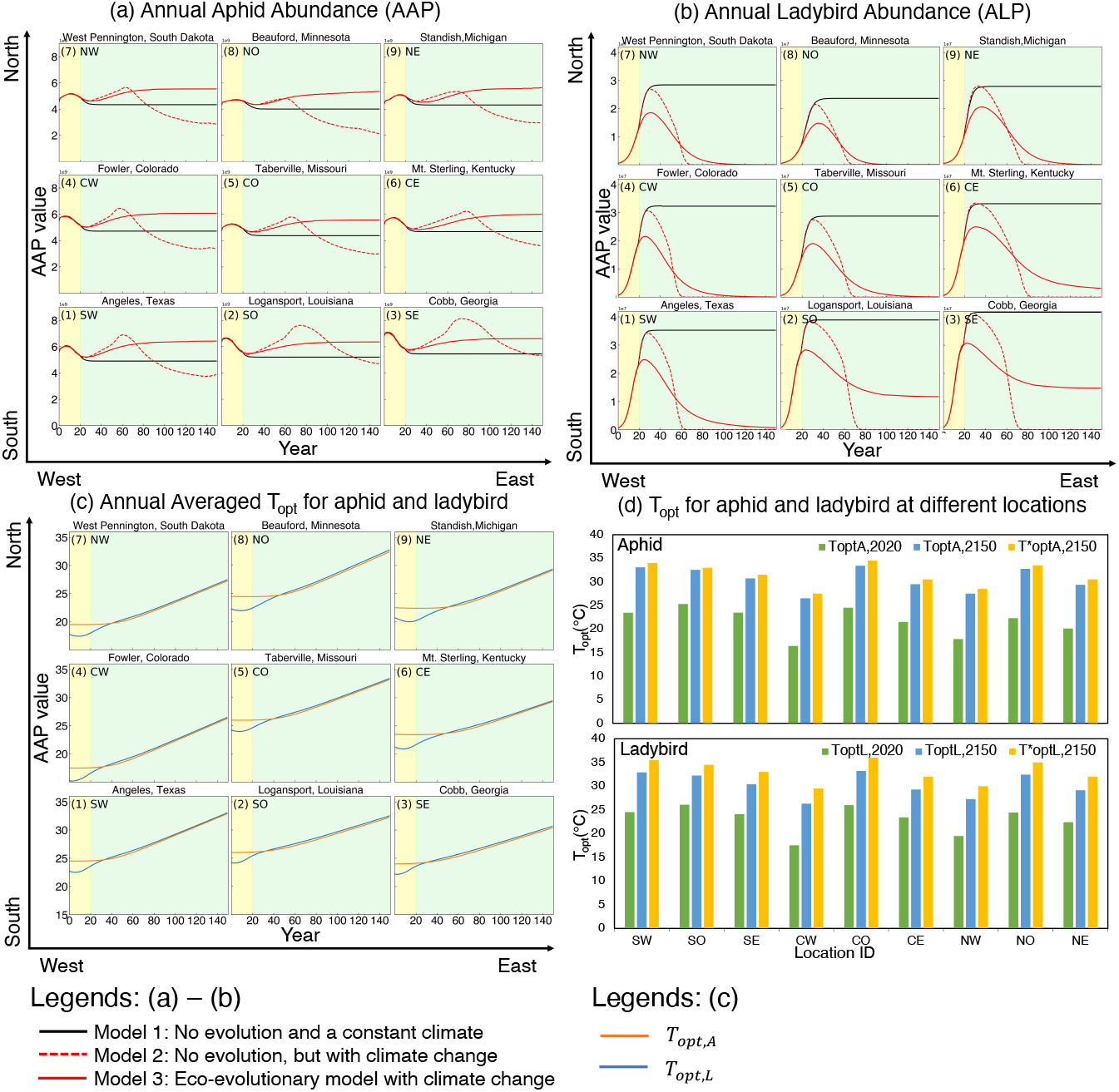
Annual population abundance and averaged *T_opt_* for aphid and ladybird over 150 years. (a) and (b) demonstrate the changing trend of annual aphid abundance and annual ladybird abundance over 150 years in different locations, each subplot demonstrates the trends for a single location. The first 20 years (yellow zone) is the flashing period for initial condition. The curves with different colors respresents the results generated by different models (see legend). (c) demonstrates the changing trend of annual averaged *T_opt_* for aphid (*T_opt,A_*, blue curve) and ladybird (*T_opt,L_*, orange curve). The barplots in (d) compares the *T_opt_* under different conditions across different locations. The green bars and blue bars represent the local adapted *T_opt,A_* and *T_opt,L_* in 2020 and 2150 generated by our eco-evolutionary model simulations. The orange bars represent the theoretical *T_opt,A_* and *T_opt,L_* which are estimated by the procedures listed in section 2.4.1.

The barplots in Fig. 4d shows the magnitudes of evolved *T_opt_* for aphid and ladybirds at different locations. The green bars represent the mean phenotypic values of *T_opt_* for aphids and ladybirds in 2020. The blue and orange bars respresent the mean phenotypic values of *T_opt_* by 2150; however, the blue bars represent the “realistic” phenotypic values estimated by the eco-evolutionary model, while the orange bars represent the “theoretical” values that match the local environment (as estimated by section 2.4.1). By comparing the phenotypic values shown in the bar plots, we can see that the evolution of aphids and ladybirds helps both species to keep pace with climate change. This is evident in the increase in their optimal thermal performance (*T_opt_*) over time. While it is not possible for evolutionary adaptation to keep up with the constantly changing climate due to the inherent time lag, evolution can still compensate for a significant portion of the effects of climate change. In terms of the ability to compensate, aphids tend to perform better than ladybirds. The bigger differences between the theoretical optimum and the evolved optimum for *T_opt,L_* compared to *T_opt,A_* indicates a lower absolute fitness for ladybirds than aphids even when *T_opt,A_* and *T_opt,L_* evolve at about the same rate (Fig. 4c). However, it is important to note that both species will still be affected by the changing climate to some extent.

### 3.2 Seasonality governs the strength of evolutionary rescue

To explore these results further, consider Logansport, Louisiana (Southern location) and Beauford, Minnesota (North-ern location). The climate in Logansport is characterized by warmer temperatures and lower seasonality compared to Beauford. Additionally, the warming trend in Logansport is slower than in Beauford. Based on these local climate con-ditions, it appears that ladybirds are able to evolve fast enough to avoid extinction in Logansport, but not in Beauford (black curves in Fig. 5 b1-b2). We explored these differences by running the model with the seasonality (s) and trend (k) parameters switched between these two locations, but leaving the baseline temperature curves (see Eq. 1) intact.

**FIGURE 5.**
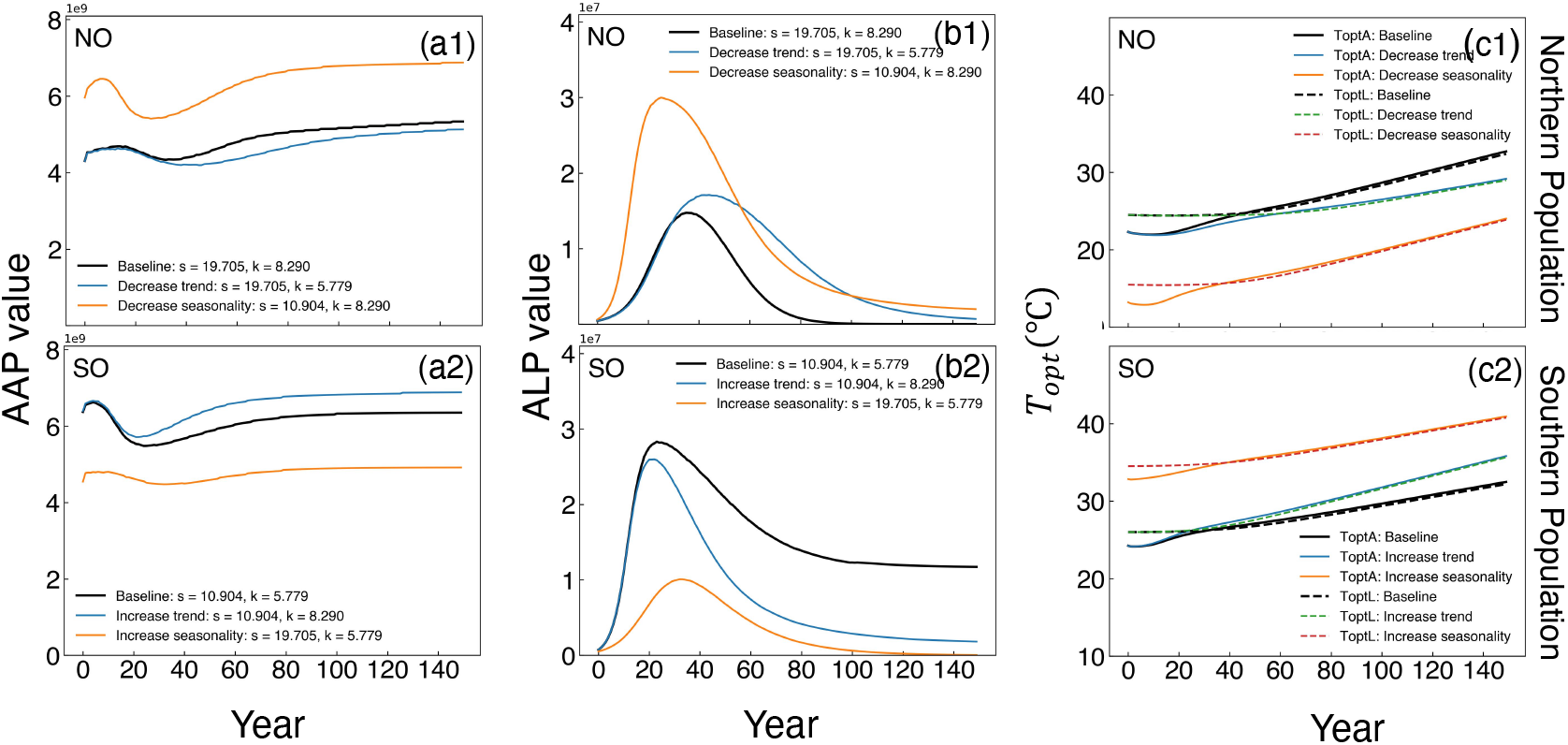
Effect of seasonality (*s*) and warming trend *k* on aphid and ladybird population abundance (*AAP* and *ALP*) and their evolving trait (*T_opt_*) in southern (SO: Logansport, Louisiana) and northern (NO: Beauford, Minnesota) locations. The blue curves represent the results generated by using the temperature parameters in each location. We exchanged the temperature parameters (*s* and *k*) between Logansport, Louisiana and Beauford, Minnesota to see which parameter is more important. Specifically, we increased the values of *s* and *k* for Logansport to the values they have in Beauford, where the climate is more seasonal and has a faster warming trend. Conversely, we decreased the values of *s* and *k* for Beauford to the values they have in Logansport. In (c1) and (c2), solid curves represent *T_opt,A_*, dashed curves represent *T_opt,L_*.

#### 3.2.1 Effects of warming trend

The warming trend and seasonality have distinct impacts on the evolution and persistence of both aphid and ladybird populations. When the warming trend in Logansport is increased, evolution still permits the ladybirds to persist, but their population abundance is lower at equilibrium (blue curve in Fig. 5 b1). When the warming trend in Beauford is decreased, ladybirds cannot avoid extinction. It simply takes longer for them to go extinct under this condition (blue curve in Fig. 5 b2). Changing the warming trend also affects the response of *T_opt_*. Faster warming tends to increase the speed of evolution for both species (Fig. 5 c1-c2).

#### 3.2.2 Effects of seasonality

The strength of evolutionary rescue in ladybirds is influenced by the seasonality of the climate. In Logansport, a more seasonal climate makes it difficult for ladybirds to adapt to climate change, leading to a decline in population size until extinction (orange curve in Fig. 5 b1). In contrast, in Beauford, evolutionary rescue occurs for ladybirds in a less seasonal climate (orange curve in Fig. 5 b2). These results suggest that the magnitude of seasonality plays a role in determining whether ladybirds are able to adapt to changing climate conditions and persist over time.

### 3.3 Effectof additive genetic variance and heritability

Figure 6 demonstrates the effect of additive genetic variance (*V_G_*) and heritability (*h*^2^) on the population abundances for aphids and ladybirds and their phenotypic values of *T_opt_*. As we see in Figs. 6c1-c2, when the additive genetic variance of aphids (*V_G_*) is high (orange curves), their *T_opt_* tends to respond a bit more rapidly (more steeply) in both southern and northern populations. However, the *T_opt_* of ladybirds remains unchanged regardless of the value of *V_G,A_.* Larger *V_G,A_* leads to decreased population abundances for both aphids and ladybirds while aphids evolve to have a higher *T_opt_*. As the additive genetic variance increases (blue-black-orange in Fig. 6a1-a2 and b1-b2), the aphid population tends to evolve to higher *T_opt_* values and incur larger intra-annual fluctuations (Fig. S1), which leads the accumulated population abundances of both aphids and ladybirds decrease.

**FIGURE 6.**
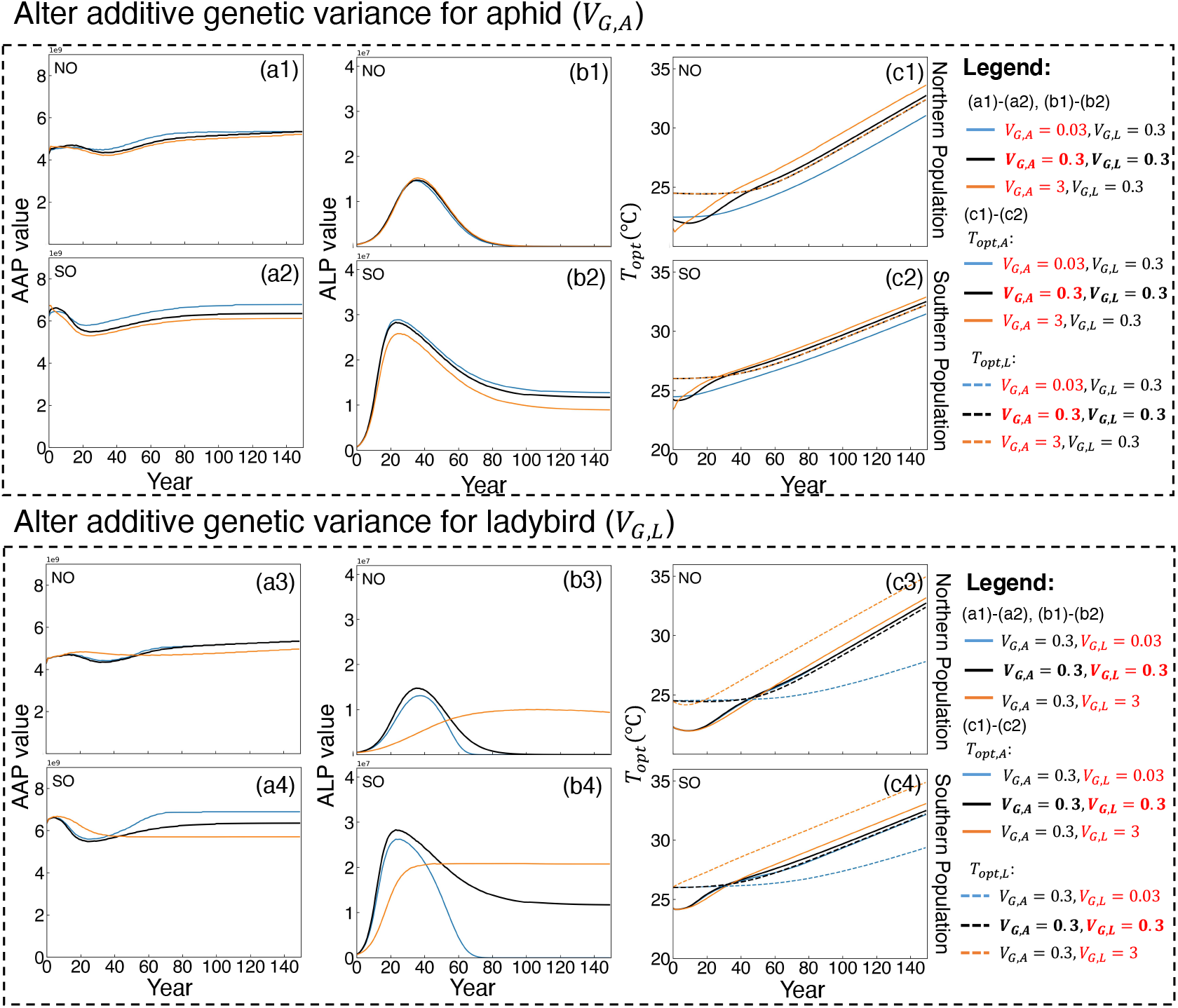
Effect of additive genetic variance and heritability on aphid and ladybird population abundance (*AAP* and *ALP*) and their evolving trait, *T_opt_*, in southern and northern locations. (a1)-(c2) represent the changing trend for *AAP*, *ALP* and *T_opt_* when altering additive genetic variance (Blue curves:V_G,A_ =0.03; Black curves:*V_G,A_* =0.3; Orange curves:*V_G,A_* =3) for aphid in southern (SO: Logansport, Louisiana) and northern (NO: Beauford, Minnesota) locations. (a3)-(c4) represent the changing trend for *AAP*, *ALP* and *T_opt_* when altering additive genetic variance (Blue curves:*V_G,L_* =0.03; Orange curves:*V_G,L_* =0.3; Green curves:*V_G,L_* =3) for ladybirds. In (c1)-(c4), solid curves represent *T_opt_*,_A_, dashed curves represent *T_opt_*,_L_.

The impact of additive genetic variance of ladybirds (*V_G,L_*) is more significant than for the aphids (*V_G,A_*). Increasing the value of *V_G,L_* (blue-black-red curves in Fig. 6c3-c4) results in stronger responses of *T_opt_* for both aphids and ladybirds, which suggests the occurrence of co-evolution. The response of ladybirds is stronger than that of aphids because *V_G,L_* directly affects the phenotype of ladybirds (Fig. 6c3-c4). Compared to *V_G,A_*. *V_G,L_* has a more significant impact on the population dynamics of both aphids and ladybirds (Fig. 6a3-a4 and b3-b4). It even plays a role in determining the persistence of the ladybird population (Fig. 6b3-b4). A larger *V_G,L_* allows ladybirds have a greater evolutionary potential to adapt to climate change and are more likely to be rescued from extinction.

Figure 7 shows the evolutionary effects on the populations of aphids and ladybirds under different evolutionary scenarios. Without the ability to adapt by selection (referred to as “evolutionary rescue”), both the southern and northern populations of ladybirds will become extinct (Fig. 7b1-b2), while the size of the aphid population will decrease in the face of climate change (Fig. 7a1-a2). If only the prey species (aphids) are able to evolve (orange curves), the aphid population will adapt to the changing environment, but the ladybirds will still be at risk of extinction, even with this indirect evolutionary effect of the aphid. If only the predator species (ladybirds) can evolve (green curve), the aphid population will not be able to adapt to the novel environment and will decrease in size. The direct evolutionary effect is still not sufficient to save the predator from local extinction. However, when both species evolve in response to climate change (black curves), evolution enables aphids to adapt to novel environments and increases the likelihood of rescuing the ladybirds from extinction (Fig. 7b1-b2). Overall, these comparisons highlight the importance of evolution effects within a predator-prey system.

**FIGURE 7.**
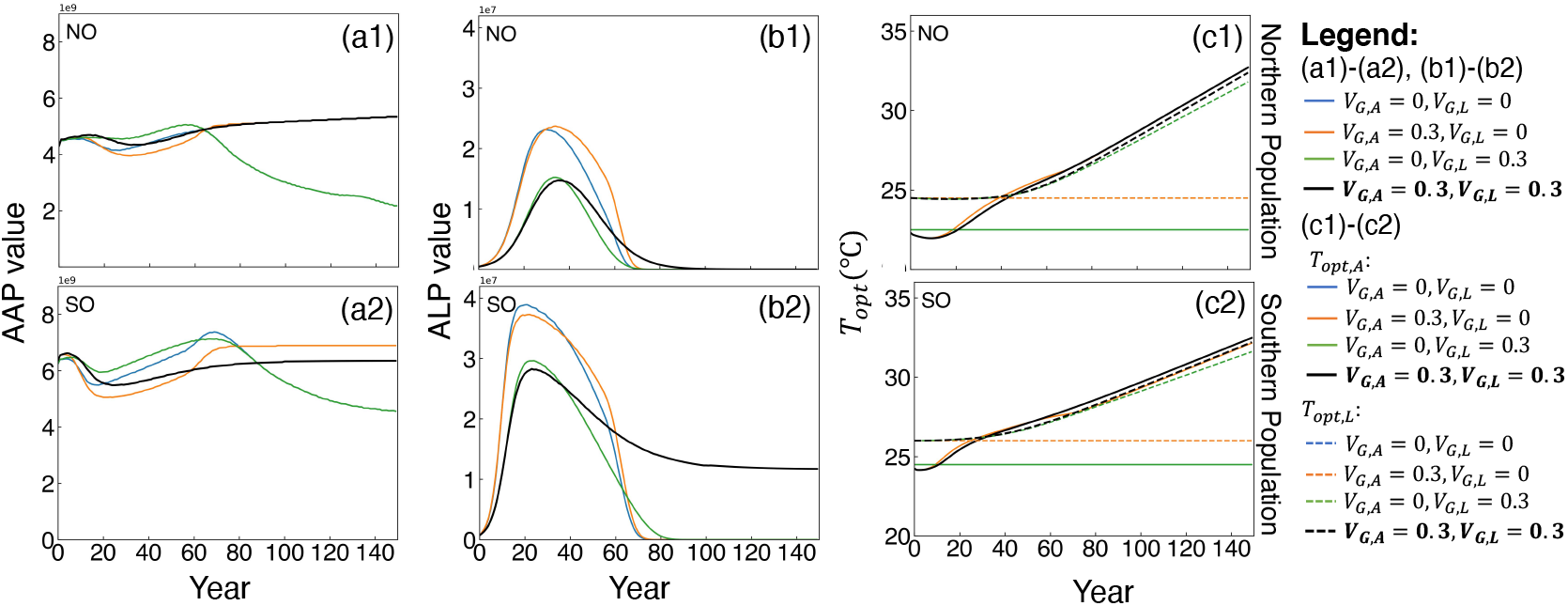
Evolutionary effects on the aphid and ladybird population abundance (AAP and *ALP*) and their evolving trait (*T_opt_*) under different evolutionary scenarios in southern (SO: Logansport, Louisiana) and northern (NO: Beauford, Minnesota) locations. Different. Blue curves (*V_G,A_* =0 and *V_G,L_*=0) represent the baseline scenario without evolution for comparison. Orange curves (*V_G,A_* =0.3 and *V_G,L_* =0) represent the evolution scenario with only prey evolves. Green curves (*V_G,A_* =0 and *V_G,L_* =0.3) represent the evolution scenario with only predator evolves. Black curves (*v_G,A_* =0.3 and *V_G,L_* =0.3) represent the scenario where aphids and ladybirds both evolve with climate change. In (c1) and (c2), solid curves represent *T_opt,A_*, dashed curves represent *T_opt,L_*.

## 4 DISCUSSION

Our eco-evolutionary model considers the interactions between different species and accounts for thermal traits that evolve in response to climate change. It utilizes the principle of marginal fitness and numerical integration to simulate changes in traits. The example of the aphid-ladybird model system demonstrates how evolutionary adaptation between interacting species affects their population dynamics over long-term climate change. Our findings suggest that there maybe geographic variation in evolutionary rescue for predators across different locations in the United States, and this rescue is primarily influenced by the change in seasonality ([*T_max,yearly_* – *T_min,yearly_*]/2). Our findings reveal that the additive genetic variance of the predator has a stronger influence on the phenotypes and population dynamics of both prey and predators compared to the additive genetic variance of the prey.

The geographical variation of evolutionary rescue seen in our study aligns with recent research that highlights the spatial patterns in evolutionary rates (Hill et al., 2011; Nadeau and Urban, 2019; Diniz-Filho et al., 2019). Our findings indicate that a predator maybe more likely to be rescued from extinction in southeastern locations. These areas tend to have a warmer, less seasonal climate with a slower rate of warming. As per our model, the potential of evolutionary rescue for locally adapted species varies across different locations based solely on the seasonality and warming trend of the location. In southeastern regions, the climates tend to be less seasonal and warming slower (Table 1). As a result, species are more able to adapt to changing conditions, thus improving their chances of survival in these areas. Diniz-Filho et al. (2019) also found a higher frequency of rescue in the eastern border of the trailing edge. Our results are similar to those of Hill et al. (2011) and Nadeau and Urban (2019), who found different evolutionary performances between the cool and warm range margins of species. Current research on the spatial patterns of evolutionary potential primarily examines individual species, leaving a need for increased study on the evolutionary dynamics of interacting species.

In our system, if only the prey evolves in response to climate change, predators will not be rescued. The “indirect evolutionary rescue” (IER) mechanism does not not seem to occur, whereby a non-evolving predator can be rescued from extinction solely due to the evolution of its prey (Yamamichi and Miner, 2015; Hermann and Becks, 2022). The IER mechanism assumes the defense cost for prey against predation relies on the predator density. If predators are scarce, prey defense is reduced, which indirectly results in an increased population growth rate of the predators (Yamamichi and Miner, 2015). Our study did not include the fitness cost of prey defense, instead focusing on the thermal effects on species vital rates. Even if the prey adapts to the changing climate and provides sufficient food resources for predators, predators may still face extinction as their thermal performances do not match the novel environment. Despite the counter-intuitive findings, our research, along with the IER mechanism, provide a theoretical basis for understanding the indirect effects of evolution.

Our findings also suggest that stronger selection for thermal performance in prey can lead to a decrease in population size, and indicates that evolutionary rescue may not always occur. Matsuda and Abrams (1994) found that the adaptive evolution of prey may ultimately lead to its own extinction by incorporating the evolution of its anti-predator ability. Such kind of process is now known as evolutionary suicide, and has been supported by other theoretical models (Gyllenberg and Parvinen, 2001; Gyllenberg et al., 2002; Vitale and Kisdi, 2019; Henriques and Osmond, 2020). Although evolutionary suicide may seem counterintuitive, the complex interplay between population dynamics and trait evolutionary dynamics may trigger the population to go extinct under certain conditions (Gyllenberg and Parvinen, 2001; Vitale and Kisdi, 2019). It would be interesting to explore the turning point at which evolutionary rescue changes to evolutionary suicide, which may provide insight into the conditions under which evolutionary rescue is more likely to occur, or the population may be at risk of extinction.

In our study, when the additive genetic variance is varied while the environmental variance is kept stable, the response of *T_opt_* aligns with Lande’s breeder equation (Lande, 1976). Bush et al. (2016) considered evolutionary adaptation only for species’ *CT_max_* and they also found that the *CT_max_* respond faster to increased heritability. These benefits to *CT_max_* asymptote at higher heritability values due to fitness costs. Our model did not account for these costs and therefore may have overestimated the evolutionary response. In addition, the magnitude of environmental effect (*V_E_*) may have uncertain consequences as an increase in *V_E_* leads to an increase in *S* but a decrease in *h*^2^. It’s not straightforward to determine whether the increase in selection differential surpasses the decrease in heritability. Therefore, it is crucial to accurately compute the selection gradient.

Most eco-evolutionary models that take into account detailed evolutionary processes are generally limited to hypothetical or simulated environments and assume simple genetics (typically one single quantitative trait) (Bocedi et al., 2014; Schiffers and Travis, 2014; Rees and Ellner, 2016). These simplifications often result in a trade-off of model uncertainty. The uncertainties in our model primarily arise from our assumptions about horizontal shifts in thermal performance and the stability of evolutionary potential (constant *V_G_*). Previous research has outlined three ways in which thermal performance curves (TPCs) can change through evolution or plasticity: (1) variation in overall performance (vertical shift); (2) variation in the thermal optimum (horizontal shift) and (3) variation in thermal specialization (Sinclair et al., 2012, their Fig. 1). These variations can also occur within geographical gradients (Tüzün and Stoks, 2018; MacLean et al., 2019). The intricate patterns in TPC variation suggest that considering different assumptions about TPCs may lead to more robust conclusions. In regards to evolutionary potential, it is commonly assumed that the additive genetic variance is constant, but some studies have found that heritability and selection of a trait can change over longer timescales (Kopp and Matuszewski, 2014; Cotto et al., 2017). Incorporating changes in evolutionary potential is a challenging task due to the lack of clear mechanisms.

Additional limitations of our eco-evolutionary model include the lack of consideration for the effects of phenotypic plasticity and dispersal. Plasticity in *T_opt_*, *CT_min_* and *CT_max_* may also play a role in driving genetic variation and can affect the rate of genetic change, either by slowing it down or accelerating it (Kopp and Matuszewski, 2014). Future research should be undertaken to incorporate additional sources of phenotypic plasticity into our eco-evolutionary model to assess the relative importance of phenotypic plasticity in evolutionary rescue. Dispersal is another mechanism which affects the evolutionary effect on species range shifts under climate change, it mitigates the negative effects of climate change and accelerates climate-induced range expansion as highly mobile individuals may be able to track suitable climates (Nadeau and Urban, 2019; Åkesson et al., 2021). While some recent research has taken into account the role of dispersal in predicting species range shifts, they have primarily focused on individual species, rather than multiple interacting species (Bush et al., 2016; Diniz-Filho et al., 2019; DeMarche et al., 2019). Future research is needed to incorporate dispersal into our eco-evolutionary model and use it as an eco-evolutionary species distribution modelling framework for interacting species.

Despite limitations, our study offers an eco-evolutionary model framework that can be incorporated into mechanistic species distribution models. Such mathematical eco-evolutionary models can provide theoretical insights into evolutionary adaptation, climate change, and biotic interactions on species population abundance and range shift.

