## Appendix S1, Figure S1, Table S1 for "Geographic variation in evolutionary rescue in a predator-prey system under climate change: an example with aphids and ladybird beetles"

### Supporting Information

**Appendix S1.** Parameters for temperature-dependent vital rates.

**Figure S1.** The daily optimum temperature for aphid and ladybird ($T_{opt,A}$ and $T_{opt,L}$) with different additive genetic variances of aphid($V_{G,A}$).

**Table S1.** Parameter values in the eco-evolutionary model.

### Appendix S1. Parameters for temperature-dependent vital rates.

Most of the parameters used in this study are roughly based on the stage-structured population dynamic model developed by Ge et al. (2022). The definitions and values for all the parameters of vital rates are listed in Table S1. As we used non-stage structured models in our study, we need to transfer the fecundity rates for aphid apterous adults and ladybird female adults and their developmental rates for pre-adult stages to the growth rate for their whole life span. We assume the growth rates for both aphids and ladybirds depend on their generation time, which is temperature-dependent. Therefore, we account for the effect of generation time on species growth rate by using the following set of equations:

$$g(\bar{z},T) = \begin{cases} m\frac{(T-CT_{min}(\bar{z}))^{q_1}(CT_{max}(\bar{z})-T)^{q_2}}{(\bar{z}-CT_{min}(\bar{z}))^{q_1}(CT_{max}(\bar{z})-\bar{z})^{q_2}} & \text{if } CT_{min}(\bar{z}) \leq T \leq CT_{max}(\bar{z}) \\ 0 & \text{otherwise} \end{cases}, \tag{A1}$$

$$GT_A(\bar{z},T) = \begin{cases} max(\frac{D_{min,A}}{g(\bar{z},T)}, D_{max,A}) & \text{if } CT_{min}(\bar{z}) \leq T \leq CT_{max}(\bar{z}) \\ 0 & \text{otherwise} \end{cases}, \tag{A2}$$

$$GT_L(\bar{z},T) = \begin{cases} max(\frac{D_{min,L}}{g(\bar{z},T)}, D_{max,L}) & \text{if } CT_{min}(\bar{z}) \leq T \leq CT_{max}(\bar{z}) \\ 0 & \text{otherwise} \end{cases}, \tag{A3}$$

$$m_A(\bar{z},T) = \frac{m_f}{GT_A(\bar{z},T)}, \tag{A4}$$

$$Q_p(\bar{z},T) = Q_p^* \times GT_L(\bar{z},T), \tag{A5}$$

where in Eq. A1, $g(\bar{z}, T)$ represents the temperature effect on developmental rate. $T$ is the air temperature. $CT_{min}$ and $CT_{max}$ are the minimum and maximum temperature thresholds beyond which developmental rate is nil. $\bar{z}$ is the mean phenotypic value of $T_{opt}$, which the optimum temperature at which the development rate reaches to its maximum. $q_1$ and $q_2$ are shape parameters which adjust the skewness of the thermal performance curve. $m = 1$, $q_1 = 1.5$, $q_2 = 1$. $CT_{min}$ and $CT_{max}$ are functions of $T_{opt}$. In Eq. A2 and Eq. A3, $GT_A(\bar{z}, T)$ and $GT_L(\bar{z}, T)$ represent the temperature-dependent generation time. $CT_{min}$ and $CT_{max}$ are the minimum and maximum temperature thresholds beyond which species couldn't develop. $D_{min,A}$, $D_{max,A}$, $D_{min,L}$, and $D_{max,L}$ are the minimum and maximum generation time for aphid and ladybird based on the experimental data. $D_{min,A} = 5$, $D_{max,A} = 20$, $D_{min,L} = 20$, and $D_{max,L} = 100$ (Satar et al., 2005; Raak-van den Berg et al., 2017). In Eq. A4, $m_A$ represents the optimal growth rate for the aphid population at optimal temperature for aphid and depends on temperature dependent generation rate. $m_f$ is the fecundity rate for aphid apterous adults, which is set as 5 per day (Xia et al., 1999; Satar et al., 2005). The optimal growth rate ($m_A$) decreases as generation time becomes longer. In Eq. A5, $Q_p$ represents the mean number of aphids a ladybird beetle offspring needs during its generation time. $Q_p^*$ is the mean number of aphids a ladybird needs to consume to reproduce a single egg, which is set as 100 aphids per day (Yu et al., 2013). $Q_p$ will become larger as generation time becomes longer.

### Supporting Figures

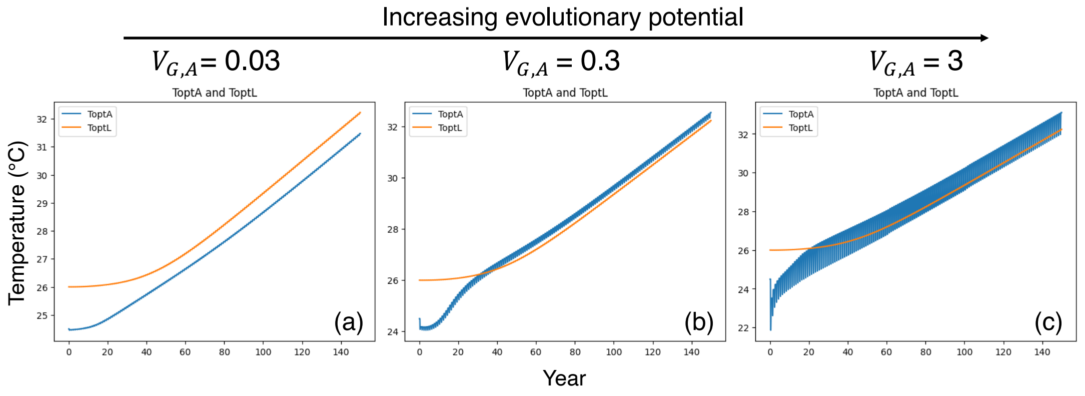


**FIGURE S1** The daily optimum temperature for aphid and ladybird ($T_{opt,A}$ and $T_{opt,L}$) with different additive genetic variances pf aphid($V_{G,A}$). As we increasing $V_{G,A}$ (a-c), the intra-annual fluctuations of $T_{opt,A}$ become larger.

**TABLE S1** Parameter values in the eco-evolutionary model.

| Rates | Parameters | Definition | Paramerer value |
|---|---|---|---|
| Aphid growth rate: $f(\bar{z}_A, T)$ | $m_A$ | Optimal growth rate at optimum temperature, its value depends on temperature-dependent generation time. | 0.25~1 |
| | $CT_{min}$ | Minimum temperature thresholds beyond which growth rate is nil, mortality rate is maximal. | $\bar{z}_A - 15$ |
| | $CT_{max}$ | Maximum temperature thresholds beyond which growth rate is nil, mortality rate is maximal. | $\bar{z}_A + 15$ |
| | $q_1, q_2$ | Shape parameters use in Eq. 12 to adjust the skewness of the thermal performance curve. | $q_1 = 1.5, q_2 = 1$ |
| Aphid intrinsic mortality rate: $\mu(\bar{z}, T)$ | $CT_{r1}, CT_{r2}$ | Temperature range in which mortality is minimal. | $CT_{r1} = \bar{z}_A - 5, CT_{r2} = \bar{z}_A + 5$ |
| | $CT_{min}$ | Minimum temperature thresholds beyond which mortality rate is maximal. | $\bar{z}_A - 15$ |
| | $CT_{max}$ | Maximum temperature thresholds beyond which mortality rate is maximal. | $\bar{z}_A + 10$ |
| | $k_1, b_1, k_2, b_2$ | Shape parameters used in Eq. 13 for aphid. | $k_1 = -0.02, b_1 = 0.45, k_2 = 0.04, b_2 = -1.15$ |
| | $\upsilon_{min}$ | Minimum mortality rate. | 0.05 |
| | $\upsilon_{max}$ | Maximum mortality rate. | 0.25 |
| Functional and numerical response | $a$ | Searching time for the ladybird to encounter an aphid. | 0.000001 |
| | $h$ | Handing time for the ladybird to process an aphid. | 0.01 |
| | $Q_p$ | Transformation rate, denotes the mean number of aphids a ladybird beetle offspring needs during its generation time, its value depends on the temperature-dependent generation time. | 2000~10000 |
| Temperature effect on functional response: $g(\bar{z}_L, T)$ | $m$ | Temperature effect on predation rate at optimum temperature | 1 |
| | $CT_{min}$ | Minimum temperature thresholds beyond which predation rate is nil. | $\bar{z}_L - 15$ |
| | $CT_{max}$ | Maximum temperature thresholds beyond which predation rate is nil. | $\bar{z}_L + 10$ |
| | $q_1, q_2$ | Shape parameters use in Eq. 12 to adjust the skewness of the temperature effect curve. | $q_1$=1.5, $q_2$=1 |
| Ladybird intrinsic mortality rate: $\gamma(\bar{z}_L, T)$ | $k_1, b_1, k_2, b_2$ | Shape parameters used in Eq. 13 for aphid. | $k_1 = -0.009, b_1 = 0.19, k_2 = 0.018, b_2 = -0.53$ |
| | $\upsilon_{min}$ | Minimum mortality rate. | 0.01 |
| | $\upsilon_{max}$ | Maximum mortality rate. | 0.1 |
